# Contrast-enhanced photon-counting micro-CT of tumor xenograft models

**DOI:** 10.1101/2024.01.03.574097

**Authors:** Mengzhou Li, Xiaodong Guo, Amit Verma, Alena Rudkouskaya, Antigone M. McKenna, Xavier Intes, Ge Wang, Margarida Barroso

**Author notes:** These authors contributed equally to this work (Mengzhou Li, Xiaodong Guo).

## Abstract

Photon-counting micro computed tomography (micro-CT) offers new potential in preclinical imaging, particularly in distinguishing materials. It becomes especially helpful when combined with contrast agents, enabling the differentiation of tumors from surrounding tissues. There are mainly two types of contrast agents in the market for micro-CT: small molecule-based and nanoparticle-based. However, despite their widespread use in liver tumor studies, there is a notable gap in research on the application of these commercially available agents for photon-counting micro-CT in breast and ovarian tumors. Herein, we explored the effectiveness of these agents in differentiating tumor xenografts from various origins (AU565, MDA-MB-231, and SKOV-3) in nude mice, using photon-counting micro-CT. Specifically, ISOVUE-370 (a small molecule-based agent) and Exitrone Nano 12000 (a nanoparticle-based agent) were investigated in this context. To improve tumor visualization, we proposed a novel color visualization method for photon-counting micro-CT, which changes color tones to highlight contrast media distribution, offering a robust alternative to traditional material decomposition methods with less computational demand. Our *in vivo* experiments confirm its effectiveness, showing distinct enhancement characteristics for each contrast agent. Qualitative and quantitative analyses suggested that Exitrone Nano 12000 provides superior vasculature enhancement and better quantitative consistency across scans, while ISOVUE-370 gives more comprehensive tumor enhancement but with a significant variance between scans due to its short blood half-time. This variability leads to high sensitivity to timing and individual differences among mice. Further, a paired t-test on mean and standard deviation values within tumor volumes showed significant differences between the AU565 and SKOV-3 tumor models with the nanoparticle-based (*p*-values < 0.02), attributable to their distinct vascularity, as confirmed by immunohistochemistry. These findings underscore the utility of photon-counting micro-CT in non-invasively assessing the morphology and anatomy of different tumor xenografts, which is crucial for tumor characterization and longitudinal monitoring of tumor development and response to treatments.

## 1. Introduction

Murine tumor models play a critical role in oncology research, enabling a wide variety of pre-clinical studies important for the diagnostic and treatment of cancer. In particular, human breast cancer lines have been widely used to generate tumor xenograft models in immunocompromised mice. These cell-line derived xenografts have been used to investigate the biology and morphology of human breast tumors (Souto *et al* 2022, Boix-Montesinos *et al* 2021). Here, we have used photon-counting contrast-enhanced micro-computed tomography (micro-CT) to characterize the anatomy and morphology of various types of breast tumor xenografts and their respective microenvironment in live, intact nude mice.

Non-invasive longitudinal studies in tumor mouse models have been performed using widely available tomographic imaging techniques, such as, magnetic resonance imaging (MRI) (Sulheim *et al* 2018, Fiordelisi *et al* 2019), optical microscopy (Smith *et al* 2022, Li 2020), ultrasound imaging (Sulheim *et al* 2018) and X-ray computed tomography (CT) (Ashton *et al* 2015, Cassol *et al* 2019). Among those modalities, X-ray micro-CT has gained significant traction for several reasons, including its noninvasive 3D imaging capability, isotropic spatial resolution (∼ 10 – 100 μm), good balance between resolution and field of view, fast speed, easy processing, and relatively low cost compared to MRI (Clark and Badea 2021). Since tumors generally have similar density to their surrounding tissues, contrast material is needed for improved tumor visibility when using micro-CT. For example, Fenestra LC is one of the most used contrast agents for liver and spleen micro-CT imaging (Bakan *et al* 2000, Sweeney *et al* 2019, Almajdub *et al* 2007, Kim *et al* 2008). Nanoparticle (NP)-based blood stream contrast agents have also been used to monitor the development of liver and spleen tumors in a longitudinal manner (Boll *et al* 2011, Cassol et al 2019, Liu *et al* 2019), via volume extraction (Cassol *et al* 2019), and vascularity (Sulheim *et al* 2018) measurements. However, compared with liver/spleen tumors, breast tumor imaging is more challenging, since it relies on positive contrast improvement based on the enhanced permeability and retention (EPR) (Maeda 2001, Wu 2021). In this context, a dedicated contrast agent with long blood half-life is desired for best vasculature tumor imaging (Sulheim *et al* 2018, Hainfeld *et al* 2018), and active targeting can be even better for specific tumor imaging (Ramesh *et al* 2022).

The micro-CT technology for small animal imaging has been rapidly evolving in recent years. Energy-integrating detectors (EIDs) are commonly used for most micro-CT scanners due to their stable performance and low cost. However, EIDs only record gray scale intensity and are often insufficient to differentiate a contrast agent from high density tissues like bones or other distinct contrast agents. Dual-energy micro-CT with dual sources was developed for material discrimination enabling imaging with two different contrast agents (Badea *et al* 2011, Clark *et al* 2013). Unfortunately, EIDs have a limited contrast sensitivity due to their higher weights for high energy photons in the image formation process. Moreover, the overlapping of the two energy spectra from the dual sources further reduces its material characterization capability. Photon-counting detectors (PCDs) have been used for micro-CT to record x-ray photons individually in an energy discriminative fashion, which allows for separation of multiple materials with superior sensitivity. Furthermore, PCDs are free of electronic noise. Given all these benefits, photon-counting CT (PCCT) can achieve similar image quality with much less radiation dose, limiting the radiation risk from multiple scans and facilitating longitudinal studies (Cassol *et al* 2019, Lu *et al* 2017). There are several commercially available photon-counting micro-CT systems for small animal imaging on the market, e.g., the MARS scanner (MARS Bioimaging Ltd., Christchurch, New Zealand) supporting simultaneous differentiation of four materials (Anderson *et al* 2010). In parallel, Siemens recently announced its first Food and Drug Administration (FDA)-approved PCCT product NAEOTOM Alpha. These technological advances have spurred the development of novel contrast agents for CT molecular imaging, benefiting tumor visualization in numerous studies. Notably, NP contrast agents based on alkaline earth metal and heavy metal (ExiTron Nano and AuroVist) have been developed and are commercially accessible, and they have gained widespread use in small animal liver CT imaging (Boll *et al* 2011, Mannheim *et al* 2016, Das *et al* 2016, Rothe *et al* 2015, Boll *et al* 2013, Cassol *et al* 2019, Mahan and Doiron 2018). Various new NP agents have also been developed in laboratory for other types of tumor CT imaging (Badea *et al* 2019, Ashton *et al* 2018, Ramesh *et al* 2022, Hainfeld *et al* 2018). However, there are few studies in which commercially available contrast agents were used for photon-counting micro-CT pre-clinical imaging of breast or ovarian tumors.

To close this gap, we have investigated two commercial contrast agents, ISOVUE-370 (human compatible iopamidol injections) and Exitrone Nano 12000 (alkaline earth metal-based NPs), for breast tumor contrast enhancement *in vivo* imaging using photon-counting micro-CT. Live nude mice carrying distinct breast and ovarian tumor xenografts were injected with different contrast agents and subjected to photon-counting micro-CT imaging. Tumor xenografts were generated from two breast cancer cells lines, which were selected as representatives of different breast cancer types: AU565 exemplifies HER2 positive breast cancer, which overexpresses epidermal growth factor 2 (HER2) receptor and lacks both estrogen (ER) and progesterone (PR) receptors and MDA-MB-231 illustrates triple-negative breast cancer lacking HER2, ER and PR receptors. SKOV-3 tumor xenografts represent HER2-positive ovarian tumors, which are well-known for their stiffness, reduced vascularity, and increased collagen content. These three tumor xenografts provide different types of tumor origin (breast vs. ovarian), receptor expression (HER2+ versus triple negative) and tumor microenvironments (high vs. low vascularity and collagen content) to demonstrate breast tumor contrast enhancement *in vivo* using photon-counting micro-CT.

Additionally, we propose a fast and convenient color visualization method to highlight the contrast media distribution without involving computationally intensive material decomposition. This color visualization method takes advantage of human eyes’ keen sensitivity to color change by correlating the spectral attenuation change with the color tone change for qualitative evaluation, and its effectiveness has been validated in our *in vivo* study. Furthermore, qualitative and quantitative enhancement effects using the two different contrast agents are compared to identify their benefits and/or disadvantages for micro-CT imaging of different types of tumor xenografts. In summary, we have investigated the potential of using photon-counting micro-CT to discriminate tumor xenografts from different origins, and cross-validated the results with immunohistochemical analysis.

## 2. Material and Methods

### 2.1 Cell culture

All human breast cancer cell lines used in this study were obtained from ATCC. HER2-overexpressing human breast cancer cell line AU565 was cultured in RPMI medium supplemented with 10% fetal bovine serum, 10 mM HEPES and 50 Units/mL/50 mg/mL penicillin and streptomycin solution. Triple negative (estrogen receptor negative, progesterone receptor negative and HER2 positive) MDA-MB-231 cells were cultured in Dulbecco’s modified Eagle’s medium supplemented with 10% fetal calf serum, 4 mM L-glutamine, 10 mM HEPES, and penicillin/streptomycin (50 Units/mL/50 mg/mL) at 37°C and 5% CO_2_. HER2-overexpressing human ovarian SKOV-3 cancer cell line was cultured in McCoy’s media, supplemented with 10% fetal bovine serum and 50 units/ml/50 µg/ml penicillin/streptomycin. All the cell lines were incubated at 37°C and 5 % CO_2_ and were used within passage ten to prevent phenotype drift and any changes in their phenotypic characteristics. Cells were routinely tested for mycoplasma to avoid contamination. For all reagents used in micro-CT imaging experiments refer to Supplementary Table 1.

### 2.2 Tumor xenografts

For generating tumor xenografts, 10 x 10^6^ AU565, 4 x 10^6^ SKOV-3, and 5 x 10^6^ MDA-MB-231 cells were mixed 1:1 with Cultrex BME and injected into the right and left inguinal mammary fat pad of 4-week-old athymic nude mice (CrTac:NCr-Foxn1nu). The subcutaneous tumors were allowed to grow for 4–5 weeks and were subjected to daily monitoring.

### 2.3 Experiments using iodine and/or NP contrast agents

To improve tumor visualization with micro-CT imaging, two commercial contrast agents, i.e., ISOVUE-370 and Exitrone Nano 12000 NPs, were injected into mice carrying tumor xenografts. To compare the two contrast agents, our experiments included three imaging sessions: mice injected with only iodine, mice injected with both iodine and NPs, and mice injected with only NPs, as detailed below. In total, 7 athymic nude mice were used in our study, and 16 effective scans have been performed (**Table 1**). At the end of each session, mice were sacrificed, and tumors were excised and fixed in formalin (3-8 ml /tumor in 15 ml tubes) for 2-5 days depending on the tumor.

1. To perform comparative imaging of different tumors using the iodine contrast agent ISOVUE-370, three athymic nude mice (tagged as AB, AL and AR) were injected subcutaneously in the right inguinal mammary pad with AU565 cells and in the opposite side with MDA-MB-231 cells. All mice had firm protruding tumors on both sides after 4-5 weeks, and sequential retroorbital injections of 120 μl iodine contrast ISOVUE-370 were performed in the anesthetized animals followed by micro-CT imaging. The injection and micro-CT imaging were repeated one week later.
2. To fairly compare the enhancement effects of different contrast agents, we injected two contrast agents into the same mice. Two athymic nude mice (tagged as JB and JR) were injected subcutaneously in the right inguinal mammary pad with AU565 cells. All mice had firm protruding tumors after 4-5 weeks, and sequential retroorbital injections of 120 μl iodine contrast ISOVUE-370 were performed in the anesthetized animals followed by micro-CT imaging. One week later, 100 ml Exitrone Nano 12000 NPs retroorbital injections were performed in the anesthetized mice and 1-2 h later, micro-CT imaging was performed.
3. To investigate the imaging enhancement effect of the NP contrast agent on different types of tumors, two athymic nude mice (tagged as S1 and S2) were injected in the right inguinal mammary pad with AU565 cells. In parallel, ovarian cancer cells SKOV-3 were injected on the left side. After protruding tumors were detected, retroorbital injections of 100 μl Exitrone Nano 12000 NPs were performed in the anesthetized animals, 1-2 h before micro-CT imaging. One week later, Exitrone Nano 12000 NPs injections followed by micro-CT were repeated. On the third week, micro-CT imaging was repeated without any additional injection.

**Table 1.** Summary of all 16 effective scans. Each mouse has a unique tag and is coded in the same color within one session. The scan tag denotes in which week the scan was performed. The combination of a mouse tag and a scan tag uniquely identifies each scan, e.g., S1-3 denotes the scan of the mouse S1 in the third week. Note for the follow-up scans S1-3 and S2-3, they were performed without contrast injection, making the time gap from last injections to current scans one week. RO = retroorbital; IP = intraperitoneal

| Session | Mouse Tag | Scan Tag | Contrast Agent | Tumor Type |  | Injection Type | Time Gap | Scan Settings |
| --- | --- | --- | --- | --- | --- | --- | --- | --- |
|  |  |  |  | Left | Right |  |  |  |
| 1<sup>st</sup> | AL | 1 | Iodine | MDA231 | AU565 | RO | <10 min | 80 kVp<br>32 μA<br>No extra filtration<br>720 views<br>250 ms |
|  | AB | 1 | Iodine | MDA231 | AU565 | RO | <10 min |  |
|  | AL | 2 | Iodine | MDA231 | AU565 | RO | <10 min |  |
|  | AR | 2 | Iodine | MDA231 | AU565 | RO | <10 min |  |
|  | AB | 2 | Iodine | MDA231 | AU565 | IP | ~ 1 hour |  |
| 2<sup>nd</sup> | JB | 1 | Iodine | NA | AU565 | RO | <10 min | 80 kVp<br>50 μA<br>1.96 mm Aluminum filtration<br>1440 views<br>300 ms |
|  | JR | 1 | Iodine | NA | AU565 | RO | <10 min |  |
|  | JB | 2 | NPs | NA | AU565 | RO | <10 min |  |
|  | JR | 2_0 | NPs | NA | AU565 | RO | <10 min |  |
|  | JR | 2 | NPs | NA | AU565 | RO | ~ 1 hour |  |
| 3<sup>rd</sup> | S1 | 1 | NPs | SKOV-3 | AU565 | RO | ~ 1 hour | 80 kVp<br>50 μA<br>1.96 mm Aluminum filtration<br>1440 views<br>300 ms |
|  | S2 | 1 | NPs | SKOV-3 | AU565 | RO | ~ 2 hour |  |
|  | S1 | 2 | NPs | SKOV-3 | AU565 | RO | ~ 2 hour |  |
|  | S2 | 2 | NPs | SKOV-3 | AU565 | RO | ~ 1 hour |  |
|  | S1 | 3 | NPs | SKOV-3 | AU565 | RO | 1 week |  |
|  | S2 | 3 | NPs | SKOV-3 | AU565 | RO | 1 week |  |

### 2.4 Immunohistochemistry

All the tumors were extracted, fixed in 10% formalin and paraffin embedded, followed by immunohistochemistry. The 5 µM tumor tissue sections were deparaffinized, rehydrated, and subjected to epitope retrieval using 1 mM EDTA pH 8.0 for 30 min. Vectastain ABC Elite kit protocol was followed for IHC staining and Vectro NovaRED was used as a peroxidase substrate. Methyl Green was used for counterstaining the tumor tissue sections. Hematoxylin & Eosin stain was used for morphological characterization and Masson’s Trichrome staining was performed for staining collagen. The tumor tissue was immunostained using HER-2 and CD 31 primary antibodies. Brightfield photomicrographs were captured using Olympus BX40 microscope equipped with Infinity 3 camera (Lumenera Inc., Ottawa, ON, Canada).

### 2.5 Photon-counting micro-CT scanning

CT scanning was performed on an adapted commercial small animal photon-counting micro-CT system (MARS16, MARS Bioimaging Ltd., Christchurch, New Zealand) with a custom-designed animal bed for anesthesia. There are 3 photon-counting chips stitched together forming a detection area of 42.24 mm x 14.08 mm, and each chip consists of 128 x 128 pixels with a pitch of 110 x 110 μm^2^. The detector supports 5 thresholds with a charge sharing correction mechanism to form 5 effective energy bins (please refer to (Getzin *et al* 2019, Jorgensen *et al* 2016) for details). During the scan, the mouse was anesthetized in a prone position with 2% isoflurane delivered via a face-cone on a customized bed for live animal imaging with its physiological status (heart rate and oxygen saturation) monitored using a paw sensor (Kent Scientific Corporation, CT, USA) clamped on its front foot as illustrated in **Figure 1**. Also, the scanner has a video camera enabling the constant monitoring of the mouse in the cabinet during imaging sessions.

**Figure 1.**
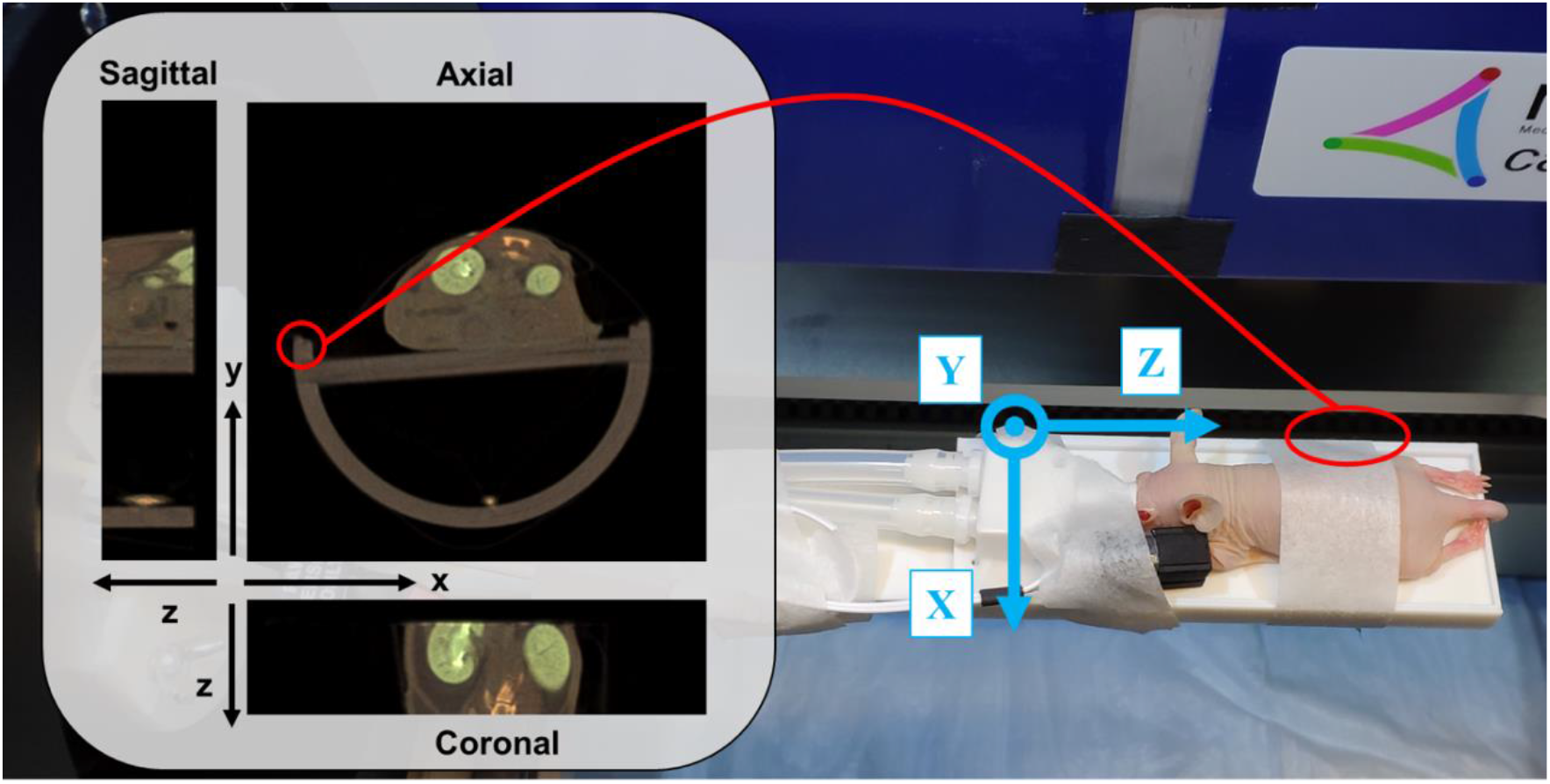
Illustration of the mouse posture and imaging setup. The associated coordinate system for image reconstruction is also displayed

To minimize the radiation dosage, we have scanned the portion of the mouse bearing tumors using a cone-beam circular scan protocol. The first imaging run was performed with the X-ray tube operated at 80 kVp, 32 μA without extra filtration to collect 720 projections per rotation with 250 ms exposure per view. To optimize the radiation dose and improve the image quality, the second and third imaging runs were performed with the tube operated at 80 kVp, 50 μA with 1.96 mm aluminum filtration to collect 1,440 projections for one rotation with 300 ms exposure time. The other parameters for all the imaging runs were kept the same. Specifically, the source to detector distance and the source to iso-center distance were 267 mm and 222 mm, respectively. X-ray photons were simultaneously counted in the energy bins of (7, 20), (20,30), (30,47), (47,73), and > 73 keV. The mouse was laid on the stomach with belly gently taped to the bed via a piece of medical surgery tape. The tumor regions were placed within the predefined field of view. The images were reconstructed at 90 μm isotropic voxel size with the MARS software. The relationship between the orientations of the reconstruction and the practical posture of the mouse is illustrated in **Figure 1**. Note that the left side of the mouse is displayed on the right of the image.

### 2.6 Summary of the micro-CT imaging scans

The effective micro-CT scans are summarized in **Table 1**. Each mouse has its unique tag and is coded in the same color within the session. The scan tag indicates which week the scan was performed, which also reflects the tumor development. Each scan could be uniquely identified with the combination of the corresponding mouse and scan tags. Hence, we will use this notation referring to one specific scan for simplicity and clarity, e.g., AL-2 refers to the scan of the mouse AL in the second week, which happened in the first session according to the name tag.

It is worth mentioning that in the first session, we performed intraperitoneal (IP) injection of iodine for the scan AB-2 and observed weak absorption of iodine into blood stream and some accumulation in urine system after one hour as shown in **Supplemental Figure 1**. The tumors were barely enhanced compared to the muscle regions as indicated by the violine/box plots of the pixel values within tumors and muscle regions, showing tumor regions do not have iodine accumulation hence posing smaller median attenuation values compared to that of the muscle region. Thus, we have sticked to the retroorbital injection method to deliver iodine into the tumor vasculature. In the second session, to illustrate the contrast enhancement in the blood stream, the liver region with big vessels and spleen regions were included in the field of view for the scan JR-2_0. JR-2 is a normal tumor scan, being different from JR-2_0.

### 2.7 Image visualization analysis

Since photon counting micro-CT provides images in multiple energy bins and different materials have their unique attenuation patterns over these bins, the material composition can be inferred from such spectral measurements for each pixel. Many material decomposition methods were proposed based on this principle, but they often require tedious system calibrations with phantoms of known material concentrations (Badea *et al* 2019, Symons *et al* 2017, Curtis and Roeder 2019, Li *et al* 2015). Furthermore, intensive computation is needed in a typical material decomposition process. For visualization of contrast-enhanced image features, these methods are widely used in a typical procedure from material decomposition, material color coding, and combination of the color maps. However, the results may suffer from high noise as the decomposition is easily influenced by the noise while the hard color coding magnifies the error visually. Hence, virtual monochromatic images are often preferred in medical practice since they are relatively insensitive to noise but require an additional computational effort. Since quantitative decomposition is not necessary in our application, to avoid the heavy computational cost, we propose a fast color visualization method to highlight the contrast agent. Taking advantage of the difference of attenuation changes between energy bins among different materials, we can highlight this kind of material associated difference to selectively highlight the used contrast agents by arranging the weighted inter-bin differences into three channels to form a RGB image. Empirically, we find that the following combinations of weights and channels achieve our goal of highlighting the iodine and NP contrast agents:

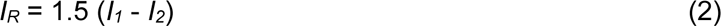

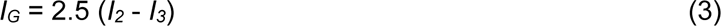

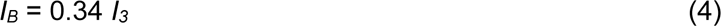

where *I_1_*, *I_2_* and *I_3_* correspond to images from bins (20,30), (30,47) and (47,73) in keV respectively, and *I_R_*, *I_G_* and *I_B_* are the three channels of the synthesized color image, respectively. The images from bins (7,20) and >73 in keV suffer from fluorescence escaping effects and high quantum noise respectively, which resulted in attenuation errors and degraded image quality, hence they were not used in this study. Note that the weights were empirically tuned for our scanning protocol, and therefore should be also adjusted for other scanning protocols or contrast agents.

### 2.8 Manual tumor annotation

The tumors were manually segmented with the software LabelMe to outline the boundaries slice by slice in either the axial or coronal view. Due to the insufficient contrast between the tumor and background, the boundaries were not always clear, producing more zigzag shapes in other views than the annotation view. As the tumor boundary should have similar smoothness in all views, directional Gaussian filtration was applied on each volume mask. Thresholding to reduce uncertainty was then implemented by leveraging the information from adjacent slices via the voting mechanism introduced by the Gaussian convolution kernel. Finally, volume masks were obtained for all the tumors presented in the micro-CT scans. It is worth mentioning that due to the limited field of view along the axial direction (Z direction), tumors were often partially covered by a circular scan not permitting adequate quantitative tumor volume measurements.

## 3. Results and Discussion

### 3.1 Color visualization

Representative images from scans with the two contrast agents are displayed in **Figure 2** using our visualization method described in **section 2.8**. From the left to right columns in **Figure 2**, the images **(a)-(d)**, **(e)-(h)**, and **(i)-(l)** correspond to scans of JR-1, JB-1, and JB-2, respectively. The first row displays the axial cross-section views, while other rows are the images from coronal views at different depths, showing the kidney, gut, and skin regions. Red boxes (axial) and circles (coronal) are used to highlight the contrast agent localized around the kidney regions, where the major differences in the enhancement effects between iodine and NP agents are revealed. Red circles in **(j)** and arrows in **(k)-(l)** point to the NP-enhanced small vessels inside the abdomen and those on the skin, whereas red arrows in **(c)-(d)** and **(g)-(h)** indicate iodine-labeled undefined vessel-like structures. Black arrow in **(g)** indicates calcium dots inside the digestive tract.

**Figure 2.**
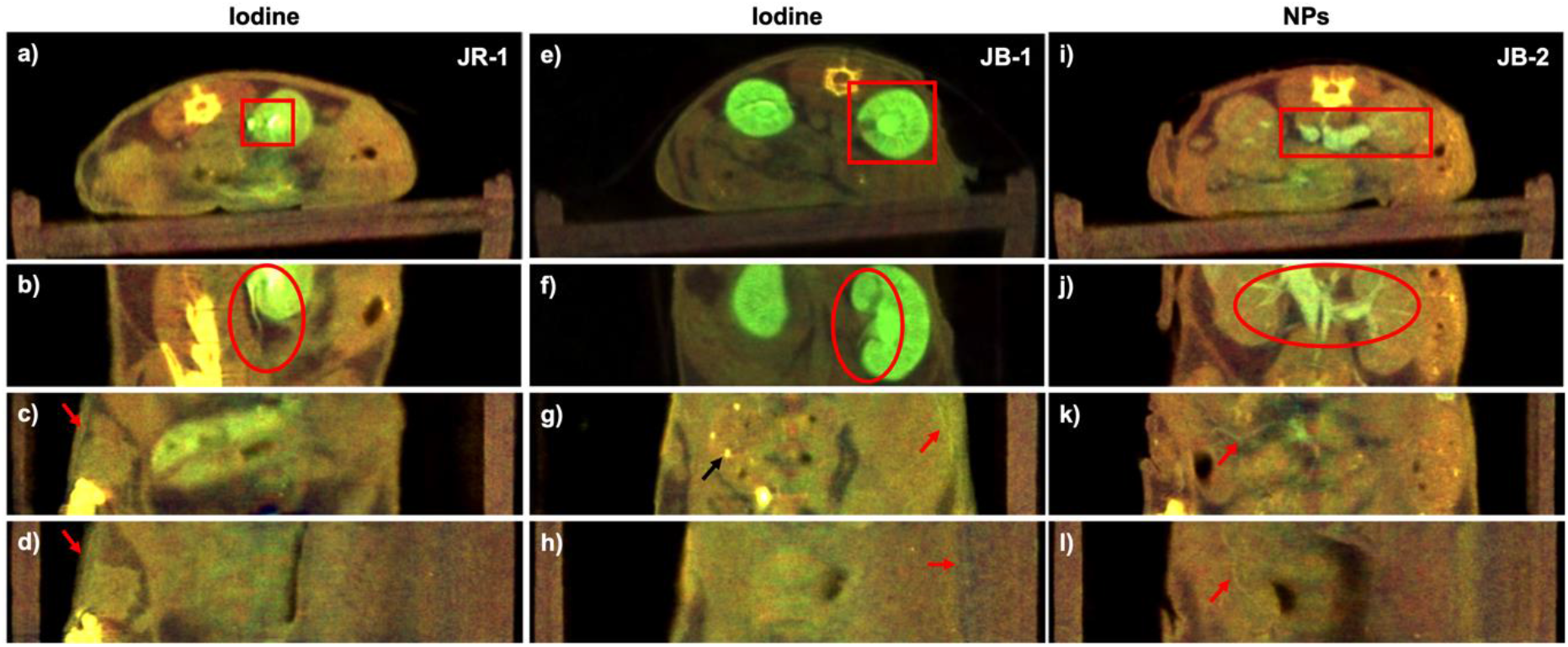
Comparison of contrast enhancement using iodine vs NP agents. **(a)-(d)**, **(e)-(h)**, and **(i)-(l)** are from photon-counting micro-CT scans of JR-1, JB-1 and JB-2 (**Table 1**), respectively. The first row shows axial images, while the remaining rows show the images from coronal views at different depths, displaying the kidney, gut, and skin regions. Yellow and brown colors represent bone and soft tissue respectively, while green color indicates the concentration of the contrast agent at the tissue location. **(e)** and **(f)** have a wider display window (doubled) than the others to avoid saturation of the high contrast details. Red boxes highlight the kidney in axial images, whereas red circles indicate the kidney vasculature in coronal views in **(b)**, **(f)** and **(j)**. NP-enhanced small vessels inside the belly and those on the skin as pointed by the red arrows in **(k)-(l)**, whereas red arrows in **(c)-(d)** and **(g)-(h)** indicate iodine labeled undefined vessel-shape structures. Black arrow in **(g)** indicates calcifications.

From the color tone, we can easily discriminate between different materials, e.g., the high intensity calcification dots in the stomach appear bright yellow in **Figure 2 (g, black arrows)** while the vessel cross-sections in the kidney appear bright green in **Figure 2 (k-l, red arrows & j, red circle)**. This analysis would be a challenge if the images were presented in gray scale with a single channel image or a single virtual monochromatic image, since they are of similar intensity levels, as shown **Figure 3 (f)**, where the bright dots inside the tumor caused by NPs are similar to the bony dots above the spine. The contrast agents concentrated in kidney or vessels were successfully highlighted in green to distinguish them from the background tissues displayed in brownish and high-density bones in yellow. Moreover, the relative concentration map of the contrast agent can be inferred from the green intensity levels, as reflected in the kidney region in **Figure 2 (b), (f) and (j)** compared to the soft tissue regions in **(c-d)**, **(g-h)** and **(k-l)**. Therefore, a qualitative material decomposition effect has been achieved using our visualization method. Since the human eye is more sensitive to the color change than to the grayscale intensity change, our method provides improved visualization and more information compared to the virtual monochromatic imaging method.

**Figure 3.**
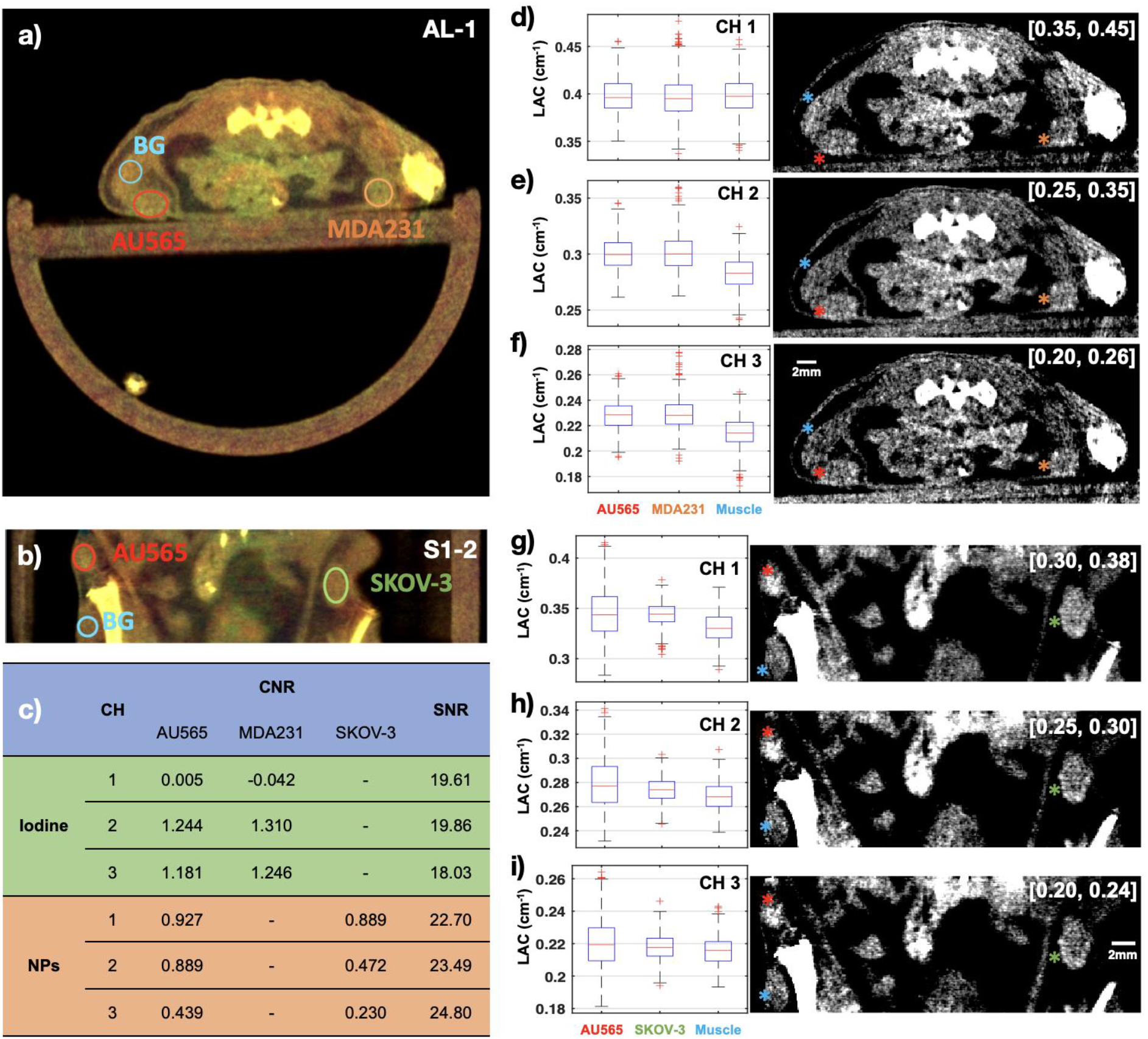
Contrast enhancement with the iodine and NP agents through visualization of exemplary slices from the axial and coronal views as shown in **(a)** and **(b)** respectively, and the corresponding gray-scale images from individual channels as shown in **(d)-(f)** for iodine, and **(g)-(i)** for NP, with display windows indicated in the right top corner of each image. The box plots of the pixel values (linear attenuation coefficient (LAC) value in unit of per cm) within tumors (AU565, MDA-MB-231, SKOV-3) and background tissues (muscle) are also displayed for each channel (CH 1-3) with the gray scale images. The contrast-to-noise ratio (CNR) with tumor as the signal and muscle as the background for each channel is tabulated in **(c)** together with the signal-to-noise ratio (SNR) calculated from the muscle regions. Right-tailed Wilcoxon rank sum tests were also performed to check the statistical significance of median value difference between tumor and muscle ROIs in each channel corresponding to **(d)-(i)**. The alternative hypothesis was accepted for all tumors that the median value of the tumor ROI is significantly greater than that of the muscle ROI with p values smaller than 0.01, except the iodine enhanced imaging in channel 1, where with the p value being 0.6165 and 0.9289 for AU565 and MDA-MB-231 the null hypothesis was accepted that the tumor data and muscle data are samples from continuous distributions with equal medians (see **Supplementary Table 2** for more details on statistical analysis).

### 3.2 Qualitative comparison between iodine and NP contrast agents

Representative results of iodine enhanced images from two scans (JR-1 and JB-1) and NP enhanced images from one scan (JB-2) are shown in **Figure 2** for comparison. The distinct appearances of the contrast-enhanced regions demonstrate that NPs are more suitable for vessel enhanced imaging, as shown in **Figure 2 (b) and (f) vs. (j), red circles**. Looking at the kidney regions marked with the red boxes or circles, the iodine contrast agent was mainly accumulated in the renal cortex, renal medulla, and renal pelvis, as well as ureter, as shown in **Figure 2 (a)-(b)** and **(e)-(f)**. Interestingly, the renal artery and renal vein appear unenhanced, such as in **Figure 2 (e)-(f)**. In contrast, injected NPs visualize the renal arteries and veins in brighter green color, while the renal cortex, renal medulla and pelvis show a much darker green tone, as shown in **Figure 2 (i)-(j)**. These differences in contrast enhancement agree well with the results shown in (Hainfeld *et al* 2018) and appear to be caused by the short half-time of the iopamidol (iodine contrast medium in Isovue-370) as well as by the ability of renal filtration to clear most of the iodine contrast agent from the blood during the 20 min scan post injection period, due to its low molecular weight. NPs that are greater than 5 nm in size do not undergo glomerular filtration, and hence they can stay longer in the blood pool (Hainfeld *et al* 2018). Based on a study with the same Exitrone 12000 NPs contrast agent described in (Cassol *et al* 2019), high metal load particles are expected to be taken up by macrophages in the liver and spleen, boosting the tissue contrast at those regions within the initial 2 h post-injection period. Similar results have been observed in our experiments as shown in **Supplementary Figure 1.** The liver and spleen tissue contrast increases continually till reaching maximum level at 24 h post-injection, with a gradual decay of tissue contrast occurring over the following 4 to 6 weeks post-injection. Consistently, with the results shown in (Cassol *et al* 2019), we detect most of the NPs in the blood stream for at least the initial several hours, allowing for the clear visualization of the big arteries and veins in **Figure 2 (i)-(j)**.

Due to NP’s longer clearing-out time from the blood pool, we can detect the NP enhanced small vessels inside the belly and those on the skin as pointed by the red arrows in **Figure 2 (k)-(l)**. However, in the iodine enhanced images, the dimmer greenish appearance all over the tissues suggests a diffused pattern for iodine within the soft tissues, as shown in **Figure 2 (c)-(d)** and **(g)-(h).** Moreover, iodine enhanced images do not display clearly identifiable vessels inside the body as shown in NP enhanced images (**Figure 2, k-l, arrows**). Although some enhanced line structures very close to skin can be seen in **Figure 2 (c)-(d)** and **(g), red arrows**, it is difficult to determine whether these structures are indeed vessels. The enhanced line between the animal skin and the bed in **Figure 2 (h)** could be urine that was excreted after iodine injection and left on the skin. Additionally, the short clearing-out time of iopamidol makes it more challenging to generate consistent image quality. As illustrated in **Figure 2**, the results of JR-1 and JB-1 gave significantly different appearances, which can be attributed to a few minutes difference in the time gaps between the injection and the scanning sessions as well as the different metabolic levels between individual mice. These small operational differences or inter-mice variance could be magnified in iodine enhanced images because of iodine’s short half-time as a contrast agent and relatively long scanning time. In contrast, the longer half-time NPs do not appear to be subjected to this image quality variability. Qualitative comparison between iodine and NP enhanced images indicates that the NP contrast agent displays improved vessel visualization. In summary, the NP contrast agent provides a high contrast enhancement in big arterials and veins, improves the visibility of small vessels, and allows for consistent image quality between scans.

### 3.3 Contrast agent effectiveness

To evaluate the effectiveness of the contrast agents, we analyzed the contrast enhancement of the tumor regions against the nearby background tissues, as shown in **Figure 3**. A typical axial slice from the scan AL-1 (AL mouse, the 1st scan) and one coronal slice from the scan S1-2 (S1 mouse, the 2nd scan) are displayed in **Figure 3 (a)-(b)**, respectively. The small regions of interest (ROI) highlighted with the red/green/orange and blue circles within tumor and non-tumor muscle tissues were selected for analysis as the tumor signal or the non-tumor background, respectively. Specifically, two metrics that are widely used in medical CT imaging evaluation (Gramer *et al* 2012), i.e., contrast-to-noise ratio (CNR) and signal-to-noise ratio (SNR), were used to quantify the contrast enhancement and the image noise levels. The CNR values were calculated by normalizing the difference in mean pixel values between the tumor ROI and the non-tumor background ROI with the standard deviation of the non-tumor background. The SNR values were calculated from only the muscle ROIs by dividing the mean value by the standard deviation. The CNR and SNR values for each channel are summarized in **Figure 3 (c)**. The SNR of the images is between 18 and 25 indicating reasonable image quality for analysis.

#### 3.3.1 Iodine enhancement of tumor xenografts

Figure 3 **(d)-(f)**, correspond to the iodine-enhanced image in **(a)**, showing the gray-scale images from individual channels and corresponding box plots of the pixel values (linear attenuation coefficient, in unit of per cm) within tumors and background tissues. The display window of each gray-scale image is indicated at the right top corner. The AU565 and MDA-MB-231 tumor ROIs are indicated by red and orange circles in **(a)** and asterisks in **(d)-(f)**, respectively, whereas background muscle ROIs are indicated by blue circles **(a-b)** and asterisks **(d-i)**. In **(e)** and **(f)** images, the red and orange asterisks appear significantly brighter than the background muscle tissue regions indicated by blue asterisk, while in **(d)** they appear similarly in brightness. The trends are evident in the respective box plots comparing the intensity distributions of those regions in three channels. The trend of channel 1 (CH 1) is different from that in the other two channels, which agrees well with channel 1 only covering the energy range right below the k-edge of iodine where the enhancement effect of Iodine is not as strong as those in channels 2 and 3 that cover the energy range above the k-edge. Our results show that more iodine agent is perfused to the tumors via vasculature and diffused to the whole tumor volume resulting in an overall increased mean value in channels 2 and 3 compared to the muscle region with strong statistical significance (*p* values smaller than 0.01 for tests on median value difference between tumor and muscle ROIs for CH2 and CH3, see **Supplementary Table 2**), as shown in Figure 3 **(e)-(f);** this can be interpreted as due to the angiogenesis and the EPR effect of tumors. Checking the quantitative values in Figure 3 **(c)**, tumor visualization is enhanced as shown by the CNR values in channels 2 and 3 (CNR > 1.0) being substantially greater than that in channel 1 (CNR ∼ 0), indicating the effectiveness of the iodine contrast agent.

#### 3.3.2 NP enhancement of tumor xenografts

Figure 3 **(g)-(i)** correspond to the NP-enhanced image in **(b)**, which displays tumors with different appearances as compared to **(d)-(f)**. For example, after iodine enhancement, AU565 tumor (red asterisks) showed a relatively uniform increased intensity across the whole tumor region **(e)-(f)**, whereas after NP injection, intensity levels across other AU565 tumors were enhanced non-uniformly **(g)-(i)**. In AU565 tumors shown in **(g)-(i)** (red asterisks), large bright spots can be detected within the tumor, most probably since the NPs cannot diffuse across the vessels easily due to their large size. Alternatively, the slice in **(b)** could be close to the surface of the AU565 tumor, a region that is richer in vessels than the center of the tumor. Brighter dots are also observed at the boundary of SKOV-3 tumors (green asterisks) than inside, while in the muscle region (blue asterisks) we did not observe such vessel aggregation pattern. Overall, NP-enhanced tumor regions are brighter than the muscle regions in all three channels, which is also supported by the box plots shown in **(g)-(i)** with strong statistical significance (*p* values smaller than 0.01 for tests on median value difference between tumor and muscle ROIs, see **Supplementary Table 2**). The NP agent appears to mainly improve the vessel contrast as NPs can hardly diffuse way from the vessels due to its large size (110 nm as indicated by the manufacturer’s specification datasheet (Miltenyi Biotec GmbH n.d.).

The NP-mediated mean value elevation and CNR trends among channels are different from those displayed by iodine-enhancement, because of the different spectral attenuation properties of NP versus iodine contrast agents. The SNR values in three channels for NP-enhancement are between 22.7 and 24.8, which are significantly higher than the values found for iodine-enhancement (SNR = 18 to 20), as shown in Figure 3 **(c)**. Such difference may be the result of different imaging settings, i.e., the combined effects of increased number of views and additional 1.96 mm aluminum filtration (see **Supplementary Figure 3**). The positive signs of all CNR values suggest the improvement of the tumor visibility in all channels, but the values for NP enhancement are smaller than those for iodine enhancement. A possible explanation for these results is that tumor vasculature only contributes partially towards an improved tumor volume visualization and therefore to increased CNR values. Alternatively, the position of the selected ROI for analysis could affect the number of vessels included. It is also worth pointing out that the images in **(g)-(i)** (NPs) might visually demonstrate a more pronounced difference between tumor and non-tumor muscle ROIs compared to panels **(d)-(f)** (iodine), which seems to contradict the CNR values in **(c)**. This discrepancy between visual perception and quantitative measurement can be attributed to two main factors. Firstly, the NP images are presented in a narrower window, resulting in a brighter appearance of the enhancement; Secondly, while the CNR measures all pixels within the tumor, NP agent only enhances the vessel pixels, which appear brighter, but the surrounding non-vessel pixels could be darker comparing to the iodine-enhanced images, thus lowering the overall CNR score. Overall, these results suggest that instead of improving the whole tumor discrimination, the NP agent mainly enhances the vessel contrast, therefore being beneficial to the vasculature imaging of tumors.

### 3.4 Tumor discrimination

Given the respiratory motion of the mouse and the resolution limit of the scanner, we are not able to directly visualize the detailed vasculature inside the tumor or perform analysis on a vascular segmentation within the tumor. Nevertheless, inspired by radiomics for medical CT, we propose to use statistical analysis of tumor pixels to reflect the vasculature information obtained from different tumors. Specifically, we have calculated the mean values and standard deviation values across the tumor volume. The mean values reflect the average amount of contrast medium in the tumor, while the standard deviation indicates the heterogeneity of the contrast medium distribution within the tumor. From a biological perspective, the former stands for the average perfusion of the tumor while the latter measures the heterogeneity of the tumor angiogenesis reflecting the distribution of effective vessels. Higher perfusion rates could lead to more contrast agent influx and higher mean values. For the diffusible contrast agent, iopamidol, higher standard deviation values could be due to the defective perfusion of tumor tissues. For example, well perfused tumors will be uniformly enhanced resulting on a smaller standard deviation in contrast to bad perfused tumors which will still have unenhanced portions due to the absence of vessels resulting on a larger standard deviation. In contrast, for the non-diffusible NP contrast agent, higher standard deviation values would indicate richer vessels (or better perfusion) since most tumor pixels would have remained unenhanced and only the vessel pixels would show increased intensity, causing the nonuniformity.

#### 3.4.1 Iodine enhancement of AU565 versus MDA-MB-231 tumor xenografts

The mean and standard deviation statistics extracted from the first imaging session (**Table 1**) are shown in Figure 4. The mean values of the two tumors followed closely in all the three channels, while AU565 always gave larger standard deviation values except for the scan AB-1. If we treat every tumor as an independent variable and plot their properties with the mean as the x coordinate and the standard deviation as the y coordinate, it is hard to separate them within the three channels. This could be caused by several reasons. First, the short clearing-out time of iopamidol could have disrupted inter-tumor relationships due to complicated combination results of inconsistencies from injection-to-imaging time intervals, metabolic level variance between mice, and different tumor characteristics. Second, the two tumor models have quite similar perfusion and vasculature characteristics and are difficult to differentiate from each other based on tumor vasculature and perfusion properties.

**Figure 4.**
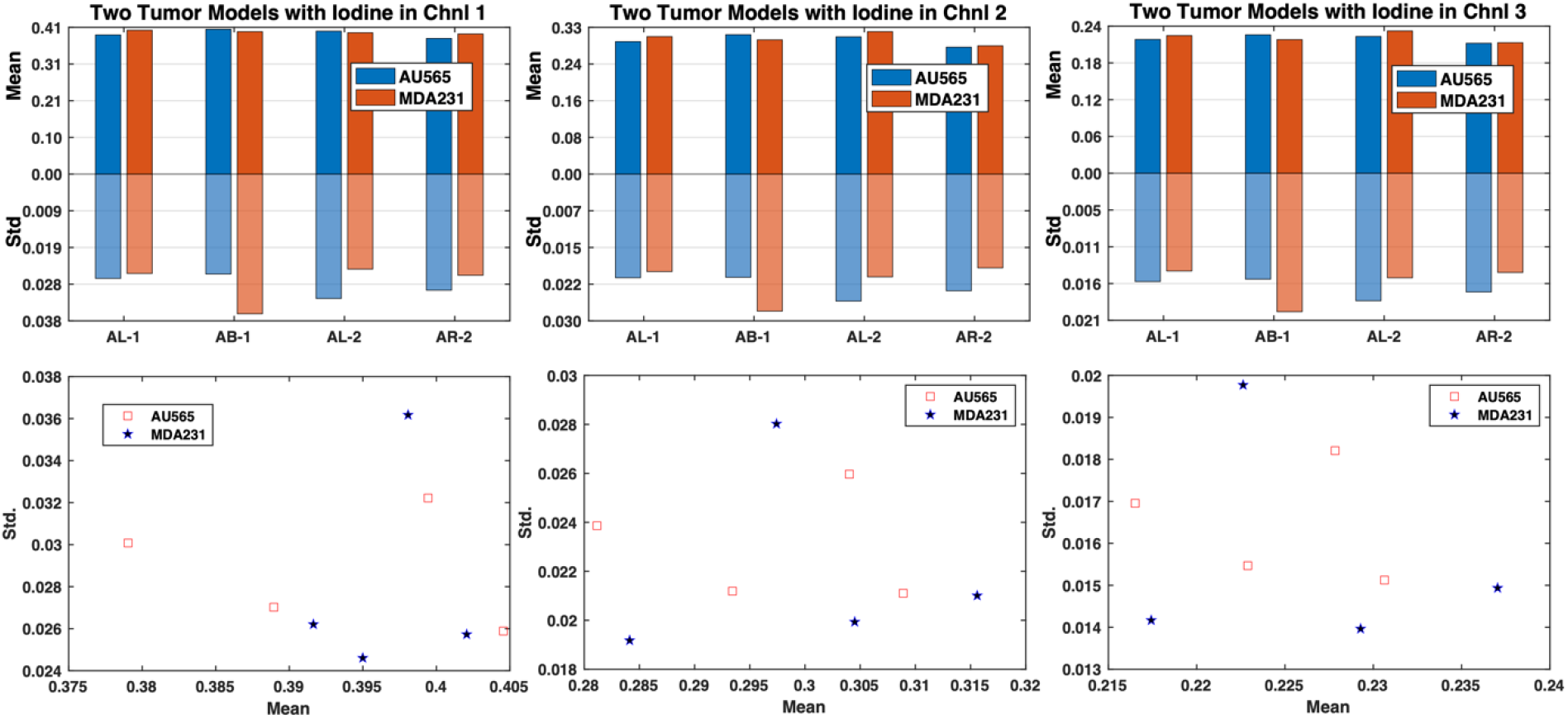
The mean and standard deviation on the AU565 and MDA-MB-231 (MDA231) tumor models from the first imaging session (AL-1, AB-1, AL-2, AR-2) using the iodine contrast agent. Top row: the bar graph comparing the mean and standard deviation of the two tumor models; Bottom row: the scatter point plot of the tumors with the mean value as the x axis and the standard deviation as the y axis.

#### 3.4.2 Nanoparticle enhancement of AU565 versus SKOV-3 tumor xenografts

The mean and standard deviation data from the third imaging session is shown in Figure 5. Different from the results extracted from the iodine imaging session, the results from the NP enhancement present consistent trends. AU565 tumors display significantly larger standard deviation and slightly greater mean values than those of SKOV-3 tumors for almost every scan and across all channels. Based on our hypothesis about the biological meaning of the mean value and the standard deviation, the results suggest that the AU565 tumors show increased perfusion and more effective vascularity than those of SKOV-3 tumors. The two tumors can be easily separated linearly in the space spanned by channels 1 and 2 when using the scatter points of the coordinates of the tumors. This suggests that the combination of mean and standard deviation values could be a metric to discriminate between AU565 and SKOV3 tumor models with the aid of the NP contrast agent.

**Figure 5.**
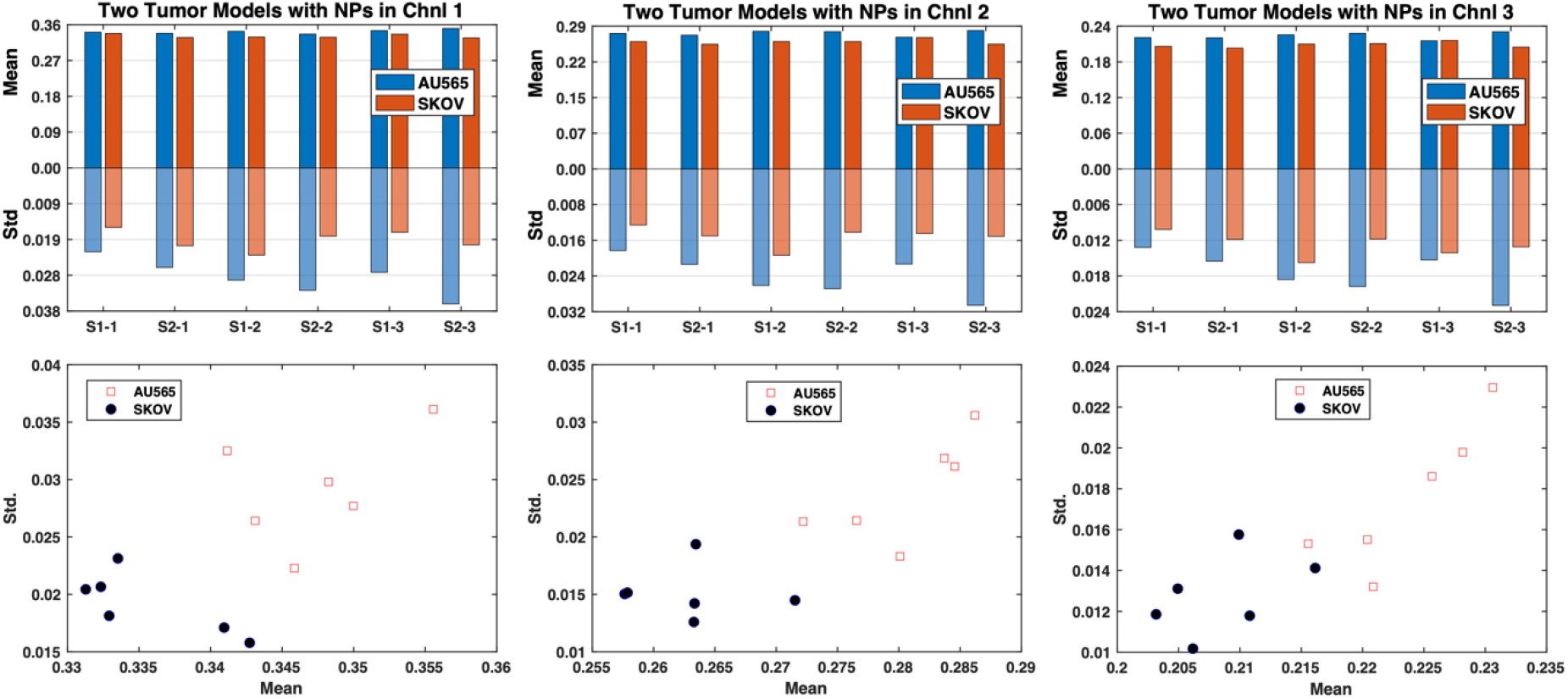
The mean and standard deviation values on the AU565 and SKOV-3 tumor models from the third imaging session (S1-1, S2-1, S1-2, S2-2, S1-3, S2-3) after the NP contrast enhancement. Top row: the bar graph comparing the mean and standard deviation of the two tumor models; Bottom row: the scatter plot of the tumors with the mean value as the x axis and the standard deviation value as the y axis.

**Figure 6.**
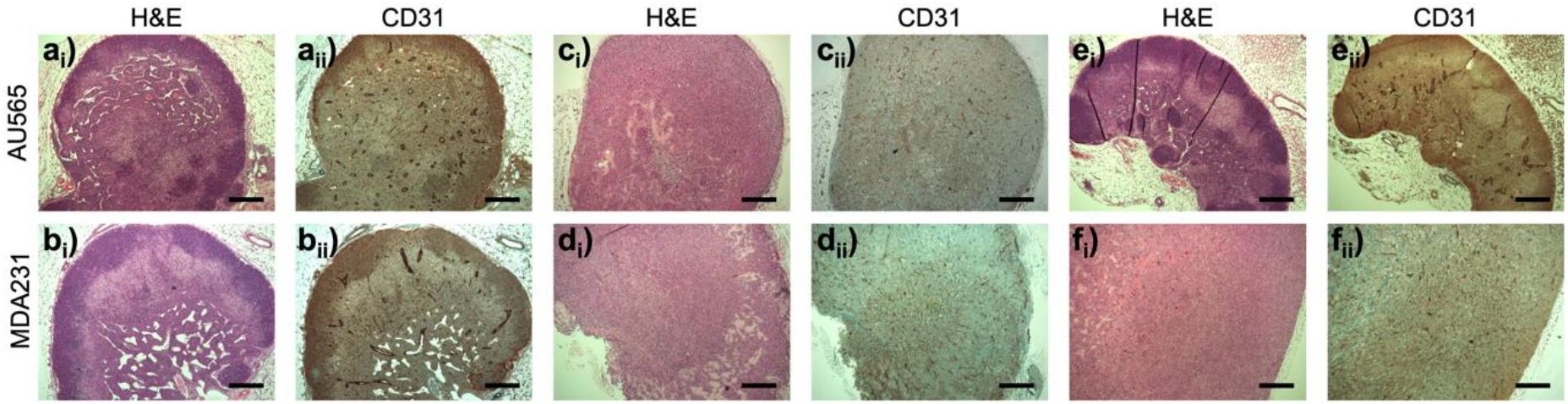
Hematoxylin and eosin (H&E) **(i)** and anti-CD31 **(ii)** staining of AU565 (**top row; a, c & e**) and MDA-MB-231 (MDA231; bottom row; **b, d & f**) tumor xenografts extracted in the first imaging session. **(a)-(b)** correspond to mouse tag AB, **(c)-(d)** mouse tag AR and **(e)-(f)** mouse tag AL (**Table 1**). Each tumor section was labeled by H&E staining **(i)**, and immunostained with anti-CD31 **(ii)** for labeling the tumor vasculature. a), b) and e) show lymph node infiltration. All the images are displayed on the same scale, and the scale bar in a) is 200 μm.**Figure 7**. Hematoxylin and eosin (H&E) **(i)**, Masson’s Trichrome **(ii)**, anti-HER2 **(iii)**, anti-CD31 staining of AU565 (**a & c**) and SKOV-3 (**b & f**) tumor xenografts extracted in the third imaging session. **(ai)-(biv)** correspond to mouse tag S1, **(ci)-(div)** mouse tag S2 (**Table 1**). Each tumor section was labeled by H&E staining **(i)**, Masson’s Trichrome **(ii)** for collagen labeling and immunostained with anti-HER2 **(iii)** or anti-CD31 **(iv)** for labeling the tumor vasculature. All the images are displayed on the same scale, and the scale bar in a) is 100 μm.

To test the statistical significance of this observation, we conducted a paired t-test on whether there was a significant difference between the mean values of the two tumors with the NP enhancement, as well as the difference between the standard deviation values of the two tumors. Since each mouse bears both tumors which were scanned simultaneously and the scan for each mouse is independent, the paired t-test was used to avoid the influence from individual difference among the mice. One-sample Kolmogorov-Smirnov test (Massey 1951) was first used to check the normality of the paired mean value differences (n=6) using the MATLAB (The MathWorks, Natick, Massachusetts) built-in function *kstest*. The normality test results suggest that we cannot reject the null hypothesis that the data came from a normal distribution, and the *p*-values for the mean value differences in three channels are 0.8871, 0.5762, 0.4807 and all significantly greater than the 5% significance level. Similar results for the standard deviation value differences were obtained with the p-values in three channels being 0.6623, 0.3178, 0.5763 and all greater than the 5% significance level (**Supplementary Table 3**). Therefore, we should accept the null hypothesis and consider that they followed normal distributions.

The paired t-test was performed as follows. Let us denote *D_i_* = *X*_1,*i*_ − *X*_2,*i*_, where *X*_1,*i*_ and *X*_2,*i*_ are the paired observations of two random variables with mean expectations being *μ*_1_ and *μ*_2_, *i* = 1, 2, ···, *n*, and *D_i_* follows a normal distribution *N* (*μ*_1_ − *μ*_2_, σ^2^). The test hypotheses are defined as:

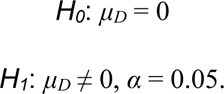

The test statistic is 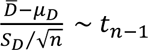, where 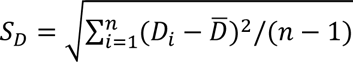. The *p*-value is calculated as 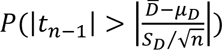.

In our case *n* = 6, and the *p*-values for mean value difference check are 0.0107, 0.0053, 0.0078 in the three channels, and similarly the p-values for standard deviation are 0.0024,0.0029 and 0.0181 in the respective channels (**Supplementary Table 4**). All are significantly smaller than 0.05, allowing the rejection of the null hypothesis at the significance level of 0.05. Thus, statistically significant differences have been established between the two tumors in terms of mean and standard deviation values with the NP contrast enhancement, indicating that AU565 and SKOV-3 tumor models can be discriminated with the aid of the NP contrast agent using micro-CT imaging, probably due to their distinct vascularity properties.

### 3.5 Immunohistochemical validation

The three representative human tumor models used in this study are MDA-MB-231, representing triple-negative breast tumors, and AU565 and SKOV-3 representing HER2 positive tumors from breast and ovarian cancer cells, respectively. analysis of MDA-MB-231 and AU565 tumor sections from the first imaging session (mouse tag: AB, AR, and AL). Qualitatively, there is no dramatic difference in the CD31 immunostaining indicating that the vasculature is similarly disrupted in both MDA-MB-231 and AU565 tumors. These results are consistent with the inability of iodine contrast agent to differentiate each tumor based on their tumor vasculature and perfusion properties using micro-CT imaging.

The immunohistochemical analysis of AU565 and SKOV-3 HER2 positive tumor sections from the third imaging session (mouse tag: S1 and S2) is shown in Figure 7. Whereas AU565 represents a non-aggressive, relatively well perfused tumor model, SKOV-3 is an example of aggressive, collagen-rich, poorly perfused tumor model (Smith *et al* 2022). Importantly, AU565, and SKOV3 HER2 positive xenografts display contrasting tumor morphologies. Although they express HER2 at similar levels, their tumor vasculature is strikingly different, with SKOV3 displaying smaller and compressed vessels, in contrast to AU565. Moreover, the collagen fiber density in SKOV3 tumors is significantly higher than that in AU565 tumors, which show increased stroma instead (Figure 7). These results correlate with the ability of NP contrast agent to discriminate AU565 from SKOV-3, probably due to their distinct vascularity properties.

**Figure 7.**
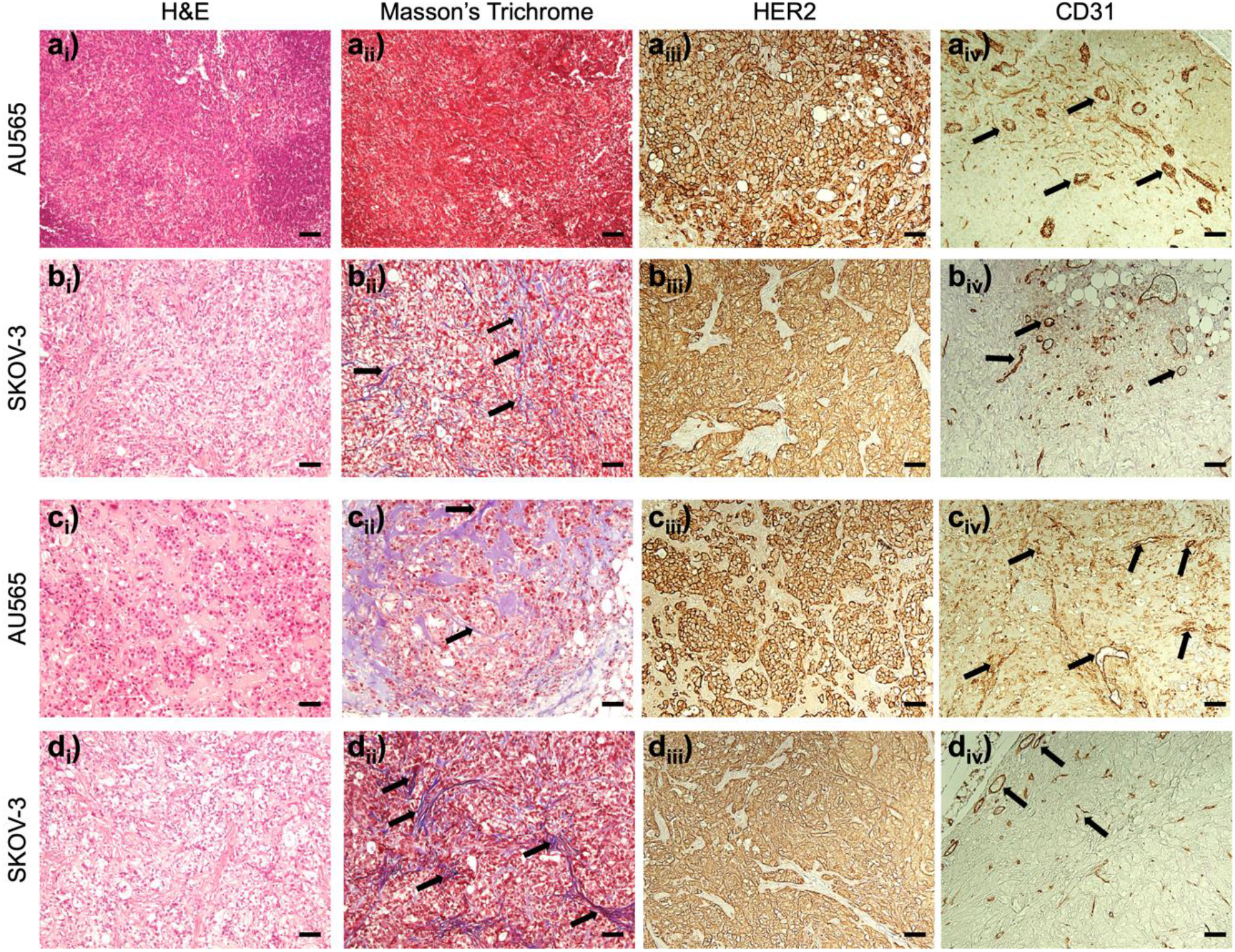
Hematoxylin and eosin (H&E) **(i)**, Masson’s Trichrome **(ii)**, anti-HER2 **(iii)**, anti-CD31 staining of AU565 (**a & c**) and SKOV-3 (**b & f**) tumor xenografts extracted in the third imaging session. **(ai)-(biv)** correspond to mouse tag S1, **(ci)-(div)** mouse tag S2 (**Table 1**). Each tumor section was labeled by H&E staining **(i)**, Masson’s Trichrome **(ii)** for collagen labeling and immunostained with anti-HER2 **(iii)** or anti-CD31 **(iv)** for labeling the tumor vasculature. All the images are displayed on the same scale, and the scale bar in a) is 100 μm.

## 4. Conclusion

In this work, we have investigated two types of commercial contrast agents, ISOVUE-370 (small molecule-based) and Exitrone Nano 12000 (NP-based), for the imaging of breast (AU565 and MDA-MB-231) and ovarian tumor xenografts (SKOV-3), for tumor contrast enhancement in live animal imaging using photon-counting micro-CT. Our proposed novel color visualization method highlights the contrast media distribution through color tone change, achieving a fast qualitative material decomposition effect that is robust to noise and easy for human to perceive. With this visualization method, two contrast agents clearly demonstrated distinct enhancement characteristics due to their large difference in cross-vessel permeability. Our qualitative comparison and quantitative analysis suggest that the nanoparticulate contrast agent provides better vasculature enhancement for each scan and better stability for quantitative analysis among scans while the iodine contrast agent gives better whole tumor enhancement but with a big variance between scans due to its short half time in blood, which means a high sensitivity to timing and individual differences among mice. Additionally, statistically significant differences in terms of mean and standard deviation values within tumors indicate that AU565 and SKOV-3 tumor models can be discriminated with the aid of the NP contrast agent using photon-counting micro-CT (*p*-values < 0.02), due to their distinct vascularity properties revealed by immunohistochemical analysis. These results demonstrate the usefulness of photon-counting micro-CT to evaluate the morphology and anatomy of distinct types of tumor xenografts non-invasively, which has a great potential in characterizing tumors and longitudinal monitoring of their development and response to treatment. On the other hand, our current results are based on scanner’s reconstruction, suffering resolution loss from mouse respiratory motion and slight geometry errors. Our next steps include upgrading the reconstruction methods to improve the resolution and SNR for direct vasculature analysis and improved tumor characterization by incorporating the latest techniques for motion compensation (Capostagno *et al* 2021, Li *et al* 2022a, 2022b), spectral distortion correction (Li *et al* 2020, Taguchi *et al* 2022), and deep learning noise reduction (Niu *et al* 2023).

## Acknowledgments

This project is funded by National Institute of Health R01 CA237267 (GW, XI and MB) and R01 CA250636 (XI and MB). We would like to thank the support of the Barroso, Intes and Wang labs.

## Ethical Statement

All animal procedures were conducted in accordance with the governing protocol approved by the Institutional Animal Care and Use Committee at both Albany Medical College (23-09001) and Rensselaer Polytechnic Institute (WANG-001-21). The animal facility at Albany Medical College has been accredited by the American Association for Accreditation for Laboratory Animals Care International.

## Data Availability Statement

The data that supports the findings of this study are available upon request.

## Author Contribution

MB, XI and GW provided critical inputs for establishing the methodology and protocols. MB and ML designed and supervised the study. ML and MB drafted and finalized the manuscript. ML and XG performed micro-CT imaging and analyzed the results. AR generated and analyzed the tumor xenografts, together with AV. AM helped with the animal imaging experiments. All authors have read, significantly revised, and approved the final manuscript.

## Supplementary data

**Supplemental Figure 1.**
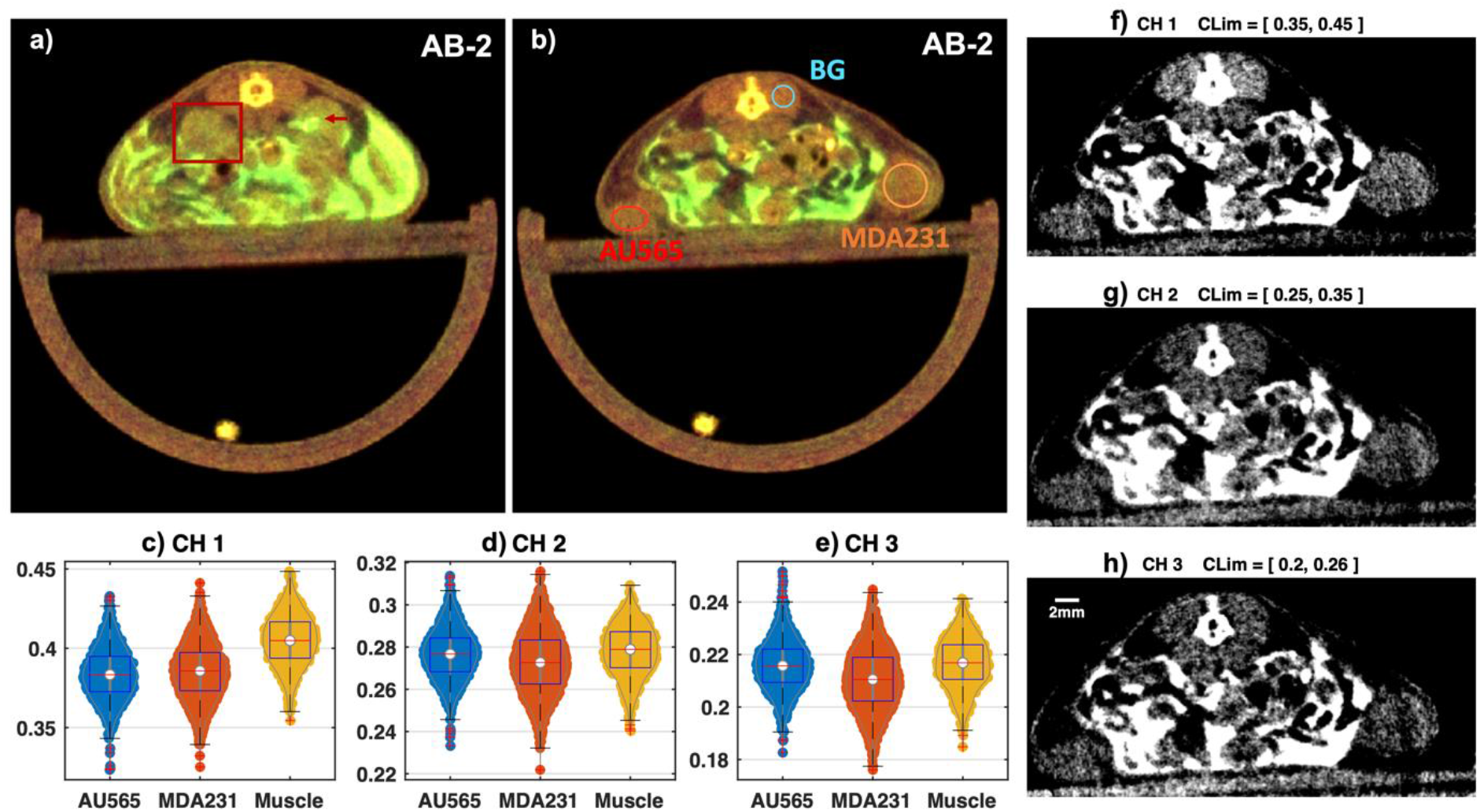
Contrast enhancement result with the iodine via IP injection (AB-2). Exemplary slices from the axial views at different depth as shown in **(a)** and **(b)** respectively. The red box and arrow in **(a)** highlight the kidney regions and the urine system showing enhanced contrast indicated by the greenish tone. The circles in **(b)** indicate the region of interest within the AU565 and SKOV-3 tumors and the muscles region (background tissue, BG) for the violin and box plots **(c)-(e)** displaying the pixel intensity values (linear attenuation coefficient, in unit of per cm) distribution for each channel. The corresponding gray-scale images for **(b)** from individual channels as shown in **(f)**, **(g)**, and **(h)**, with display windows indicated at the top of each image.

**Supplemental Figure 2.**
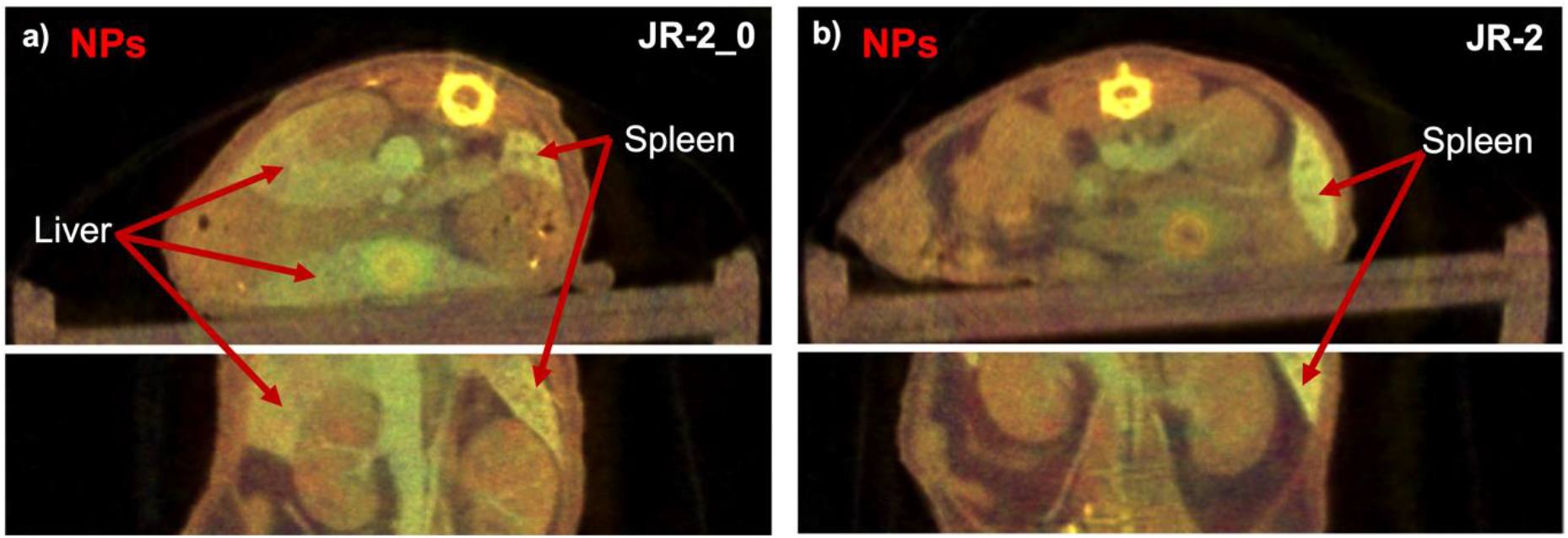
Contrast enhancement results with NPs show strong accumulation of contrast agents in liver and spleen. Exemplary slices from scan JR-2_0 and JR-2 are shown in **(a)** and **(b)** respectively, with axial views in the top row and coronal views in the bottom row. The liver and spleen regions are indicated by red arrows. Note that JR-2_0 was 1 hours later than JR-2, hence, the spleen regions in JR-2 appear brighter than those in JR-2_0 due to the gradual taken-up of NPs by macrophages (Cassol *et al* 2019).

**Supplemental Figure 3.**
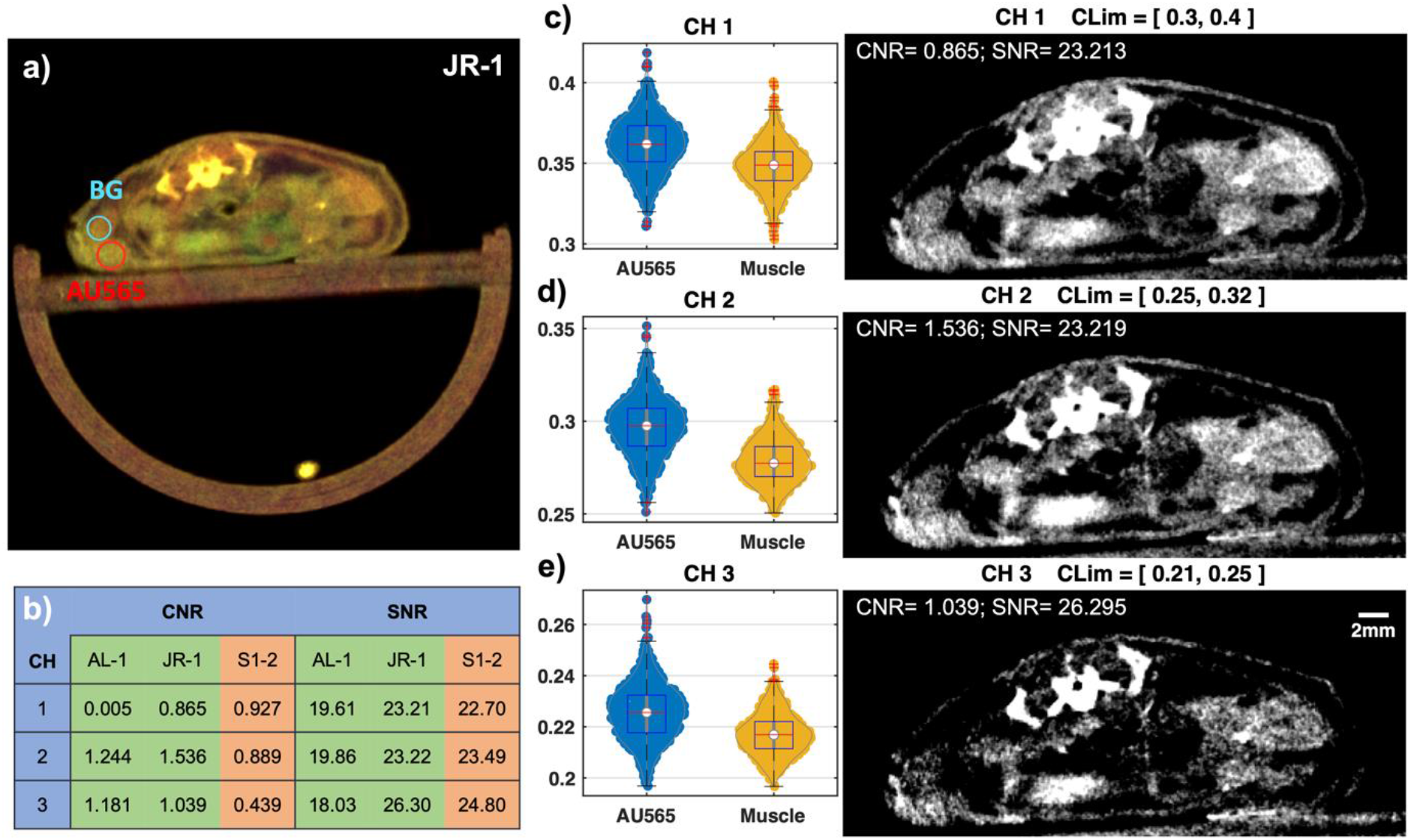
Iodine contrast enhancement results with the same imaging settings as that of NPs enhanced scans. Visualization of one axial slice from the scan JR-1 is shown in **(a)**, and the corresponding gray-scale images from individual channels are shown in **(c)-(e)** with display windows indicated at the top of each image. The violin and box plots of the pixel values within the tumor (AU565) and background tissues (muscle) are also displayed for each channel (CH 1-3) with the gray scale images. The CNR values for AU565 and SNR values in each channel are tabulated in **(b)** together with the values from the other scans demonstrated in **Figure 3**. JR-1 share similar SNR values with S1-2 but not with AL-1, suggesting the SNR discrepancy between AL-1 and JR-1 is mainly caused by their different imaging settings. The CNR pattern of JR-1 is similar to that of AL-1 in channels 2 and 3, which may attribute to the iodine imaging characteristic. The major difference in channel 1 should result from the additional aluminum filtration altering the X-ray spectrum for imaging, e.g., significantly reducing low energy photons that fall within the channel 1.

**Supplementary Table 1.** Reagents used in micro-CT imaging experiments.

| **Supplementary table 1. Reagents used in micro-CT imaging experiments** |  |  |
| --- | --- | --- |
| Resources | Source | Catalog number |
| Nude mice, CrTac: NCr-Foxn1nu | Taconic Biosciences, Rensselaer, NY | NCRNU-F |
| Exitrone nano 12000, Alkaline earth metal-based NPs | Miltenyi Biotec GmbH, Germany | 130 - 095 - 698 |
| ISOVUE-370 | Human compatible iopamidol injections, Bracco, Milan, Italy | CL67E08 |
| HER2/ErbB2 (29D8) Rabbit mAb | Cell Signaling, Technology, Danvers, MA | 2165 |
| Monoclonal rabbit CD31 | Cell Signaling Technology, Inc., Danvers, MA | 77699 |
| Vectastain ABC Elite kit | Vector Labs, Burlingame, CA | PK-6100 |
| Vectro NovaRED | Vector Labs, Burlingame, CA | SK-4800 |
| Methyl Green | Sigma Aldrich, St. Louis, MO | M8884 |
| AU565 | ATCC, Manassa, VA | CRL-2351 |
| SKOV-3 | ATCC, Manassa, VA | HTB-77 |
| MDA-MB-231 | ATCC, Manassa, VA | HTB-26 |
| RPMI 1640 | Thermo Fisher Scientific Inc, Waltham, MA | 11875093 |
| McCoy's media | Thermo Fisher Scientific Inc, Waltham, MA | 16600082 |
| Fetal bovine serum | ATCC, Manassa, VA | 30-2021 |
| HEPES | Thermo Fisher Scientific Inc, Waltham, MA | 15630080 |
| L-Glutamine | Thermo Fisher Scientific Inc, Waltham, MA | 25030081 |
| BME001-05 | R&D Systems Inc, Minneapolis, MN, USA | BME001-01 |

**Supplementary Table 2.** p-values of the statistical tests on the comparison of median values between tumor ROIs and muscle background ROIs (BG) in Figure 3. Since the pixel value distribution does not necessarily follow a normal distribution for each ROI, e.g., the MDA-MB-231 (MDA231) at channel 2 does not pass the one-sample Kolmogorov-Smirnov test, a right-tailed Wilcoxon rank sum test rather than t-test was employed to assess any significant difference between the median values of ROIs from tumor (x) and muscle regions (y) across each channel corresponding to Figure 3 (d)-(i). The null hypothesis is that the tumor data and muscle data are samples from continuous distributions with equal medians, while the alternative hypothesis is that the median of tumor data is greater than that of muscle data. For iodine contrast imaging, the median value of both tumor regions are significantly greater than the median value of muscle regions in channels 2 and 3 (p<0.05), while we cannot reject the null hypothesis for the channel 1 with p values being 0.6165 and 0.9289 for AU565 and MDA231. For NPs contrast imaging, with all p values being smaller than 0.05, we can conclude that the median values of both AU565 and SKOV-3 tumors are statistically greater than that of muscle regions in all three channels. * Note the p value gets zeros due to limitations of numerical precision in computing. The test was performed using the MATLAB built-in function *ranksum*. For statistical analysis, the ROIs contained 753 points for AU565, 609 for MDA231, and 421 for muscle background in the iodine enhanced imaging. In NP-enhanced imaging, the ROIs contained 455 points for AU565, 665 for MDA231, and 305 for muscle background.

| CH | Iodine |  | NPs |  |
| --- | --- | --- | --- | --- |
|  | AU565 v.s. BG | MDA231 v.s. BG | AU565 v.s. BG | SKOV-3 v.s. BG |
| 1 | 0.6165 | 0.9289 | 0.0000* | 0.0000* |
| 2 | 0.0000* | 0.0000* | 0.0000* | 0.0000* |
| 3 | 0.0000* | 0.0000* | 0.0001 | 0.0002 |

**Supplementary Table 3.** p-values of the statistical tests on whether the paired value differences among six scans follow a normal distribution in terms of mean and standard deviation values for each channel (Figure 5). The null hypothesis is that pair differences (six samples) between the AU565 tumor data and SKOV-3 tumor data are samples from a normal distribution while the alternative hypothesis is that they do not come from a normal distribution. The difference values are normalized to zero mean and unit variance and then fed to the MATLAB built-in function *kstest* for the statistical test. All *p*-values in both terms of mean differences and standard deviation differences are significantly greater than the 0.05 threshold for all three channels. Thus, we should accept the null hypothesis that those values follow a normal distribution, making they eligible for a paired *t*-test followed.

| CH | Paired value differences between AU565 and SKOV-3 among 6 scans after normalization |  |
| --- | --- | --- |
|  | Mean value differences | Standard deviation differences |
| 1 | 0.8871 | 0.6623 |
| 2 | 0.5762 | 0.3178 |
| 3 | 0.4807 | 0.5763 |

**Supplementary Table 4.** p-values of the paired *t*-tests on whether there is a significant difference between the samples from AU565 and SKOV-3 among six scans in terms of the mean values and standard deviations of the tumors for each channel (Figure 5). The null hypothesis is that the means of two paired populations respectively from the AU565 tumor and SKOV-3 tumor are equal while the alternative hypothesis is that they are not equal. All *p*-values in both terms of mean values and standard deviation values are smaller than the 0.05 threshold for all three channels. Thus, we can reject the null hypothesis that the mean values or standard deviation values for the AU565 and SKOV tumors are the same.

| CH | Paired value from AU565 and SKOV-3 among 6 scans |  |
| --- | --- | --- |
|  | Test on mean values | Test on standard deviation values |
| 1 | 0.0107 | 0.0024 |
| 2 | 0.0053 | 0.0029 |
| 3 | 0.0078 | 0.0181 |

